# Neurophysiological Changes Associated with Vibroacoustically-augmented Breath-focused Mindfulness for Dissociation: Targeting Interoception and Attention

**DOI:** 10.1101/2023.01.04.521609

**Authors:** Negar Fani, Alfonsina Guelfo, Dominique L. La Barrie, Andrew P. Teer, Cherita Clendinen, Leyla Karimzadeh, Jahnvi Jain, Timothy D. Ely, Abigail Powers, Nadine Kaslow, Bekh Bradley, Greg J. Siegle

## Abstract

**Background:** Dissociative symptoms can emerge after trauma and interfere with attentional control and interoception, both of which are barriers to mind-body interventions such as breath-focused mindfulness (BFM). To overcome these barriers, we tested the use of an exteroceptive augmentation to BFM, using vibrations equivalent to the amplitude of the auditory waveform of the actual breath, delivered via a wearable subwoofer in real time (VBFM). We tested whether this device enhanced interoceptive processes, attentional control and autonomic regulation in trauma-exposed women with dissociative symptoms.

**Method:** 65 women, majority (82%) Black American, aged 18-65 completed self-report measures of interoception and 6 BFM sessions, during which electrocardiographic recordings were taken to derive high-frequency heart rate variability (HRV) estimates. A subset (n=31) of participants completed functional MRI at pre- and post-intervention, during which they were administered an affective attentional control task.

**Results:** Compared to those who received BFM only, women who received VBFM demonstrated greater increases in interoception, particularly their ability to trust body signals, increased sustained attention, and as well as increased connectivity between nodes of emotion processing and interoceptive networks. Intervention condition moderated the relationship between interoception change and dissociation change, as well as the relationship between dissociation and HRV change.

**Conclusions:** Vibration feedback during breath focus yielded greater improvements in interoception, sustained attention and increased connectivity of emotion processing and interoceptive networks. Augmenting BFM with vibration appears to have considerable effects on interoception, attention and autonomic regulation; it could be used as a monotherapy or to address trauma treatment barriers.

---

Some survivors of psychological trauma develop symptoms of dissociation, which is characterized by feelings of detachment from the self (depersonalization) and/or surroundings (derealization). Some estimates indicate that 15-30% of trauma survivors with posttraumatic stress disorder (PTSD) exhibit clinically significant dissociation(Stein et al., 2013; Wolf et al., 2012). Particularly when faced with trauma-relevant information, these individuals become highly distressed, experience involuntary disengagement of attentional focus, and disconnect from their bodies(Zerubavel & Messman-Moore, 2015). This may impede therapeutic engagement, particularly for trauma-focused treatments; in fact, dissociative symptoms have been linked to poor response to first-line treatments such as prolonged exposure therapy(Jaycox, Foa, & Morral, 1998; Kleindienst et al., 2011; Michelson, June, Vives, Testa, & Marchione, 1998; Spitzer, Barnow, Freyberger, & Grabe, 2007). For dissociative trauma-exposed individuals, increased distress in the presence of trauma cues has also been correlated with autonomic dysregulation, in the form of blunted heart rate reactivity(Griffin, Resick, & Mechanic, 1997; Lanius et al., 2002) and low heart rate variability (HRV)(Reinders et al., 2006); HRV references the variability within the beat-to-beat interval, and is thought to index cardiac vagal tone, or vagally-mediated parasympathetic activity(Laborde, Mosley, & Thayer, 2017).

Given that dissociative states involve detachment from the body and/or surroundings, this naturally hinders interoception, a term that references a set of functions related to the ability to detect, accurately interpret, and trust internally-generated physiological signals(Schaflein, Sattel, Pollatos, & Sack, 2018). Interoception provides a basis for successful emotion regulation(Price & Hooven, 2018); identifying physiological signatures of reactivity allows individuals to choose how to modulate these reactions. Detachment from the self and/or surroundings is a state that is incompatible with voluntary control of attention and interoceptive awareness, which is apparent from brain and behavior studies; dissociative trauma-exposed individuals demonstrate disruptions in interoceptive brain networks and concomitant dysregulation of attention in the context of emotion(Fani et al., 2018; Krause-Utz, Frost, Winter, & Elzinga, 2017). Increasingly, abnormalities in interoceptive functions are thought to play a role in stress-related disorders(Schulz & Vogele, 2015) including clinically-significant dissociation(Dyer, Feldmann, & Borgmann, 2015; Michal et al., 2014; Pick et al., 2020; Schaflein et al., 2018; Schulz et al., 2015; Sedeno et al., 2014).

Mindfulness-based interventions (MBIs) may benefit dissociative tendencies(Boyd, Lanius, & McKinnon, 2018; Didonna et al., 2019; Price, Wells, Donovan, & Rue, 2012; Sharma, Sinha, & Sayeed, 2016), possibly via effects on interoceptive awareness. MBIs involve sustained, non-judgmental present-centered attention, often to sensory cues such as the breath. Given that attention to sensory cues is targeted in MBIs, these practices may be well-suited to address interoceptive deficits in dissociative trauma-exposed people, which may, in turn lead to decreases in dissociative symptoms. Increases in self-reported interoceptive awareness have been observed following MBIs(Bornemann, Herbert, Mehling, & Singer, 2014; de Jong et al., 2016; Fissler et al., 2016). This may be secondary to changes in neural networks that support interoception, attentional control and emotion regulation; some evidence suggests that MBIs lead to increased interoceptive network connectivity and response in fronto-parietal attentional control neural networks(Farb, Segal, & Anderson, 2013; Goldin, Ziv, Jazaieri, & Gross, 2012). However, there is little information on how changes in self-reported interoceptive awareness following mindfulness treatment relate to neurophysiological alterations as well as changes in dissociation.

Emerging data suggests that interoceptive awareness can be improved with exteroceptive signaling. Exteroceptive and interoceptive signals interact to inform a person’s sense of self(Salvato, Richter, Sedeno, Bottini, & Paulesu, 2020); exteroceptive information can affect interoception, including feelings of self-agency and body trusting. For example, the classic “rubber hand illusion” experiment demonstrated that the timing of exteroceptive (visual and tactile) feedback influences sense of body ownership(Botvinick & Cohen, 1998). Notably, dissociative individuals with PTSD demonstrate a stronger illusion effect as compared to non-dissociative individuals, showing difficulties in differentiating real and imagined touch and simultaneously affecting escalation in dissociative symptoms during the experiment(Rabellino et al., 2018). Some researchers have observed that providing exteroceptive feedback with cardiac signals *enhances* feelings of body ownership and interoceptive sensitivity(Suzuki, Garfinkel, Critchley, & Seth, 2013). Further research indicates that exteroceptive signals serve to trigger the integration of exteroceptive and interoceptive sensations in the insula, which serves as a hub for interoception(Koeppel, Ruser, Kitzler, Hummel, & Croy, 2020). As such, exteroceptive cues may be grounding stimuli that can penetrate dissociative states and enhance interoceptive awareness(Ogden, Pain, & Fisher, 2006). However, no studies to date have tested the use of exteroceptive feedback during MBIs with dissociative individuals.

We examined the effects of exteroceptive augmentation of an MBI, breath-focused mindfulness, on interoceptive processes and attention to emotion in a sample of dissociative, trauma-exposed individuals, a majority of whom were Black American. As an exteroceptive feedback modality, we tested an experimental intervention that incorporated vibrations equivalent to the amplitude of the auditory waveform of individuals’ actual breath in real time (“vibroacoustic stimulation”), with vibrations delivered via a small subwoofer worn on the chest. We used an affective attentional control task (Affective Stroop, AS) during functional MRI to assay potential changes in amygdala connectivity to other emotion processing and interoceptive network regions. Participants were randomly assigned to one of two breath-focused mindfulness (BFM) intervention conditions, one of which involved vibroacoustic augmentation during BFM (VBFM) delivered via a device placed on the sternum; the BFM group wore the device but received no feedback.

All participants underwent six 15-minute sessions of their assigned intervention condition with concurrent electrophysiology recording. A subset of participants completed neuropsychological testing (n=42) and received the AS (n=31) during fMRI at pre- and post-intervention. Interoceptive awareness was assessed with a validated measure that assesses different interoception dimensions(Mehling et al., 2012). We hypothesized that those who received VBFM would experience greater increases in interoception and improvements in attentional control (assayed via neuropsychological tests) as compared to BFM without exteroceptive augmentation. We also hypothesized that vibration feedback would moderate the relationship between change in interoception and change in dissociation, as well as change in two physiological mechanisms: autonomic regulation (HRV) and neural network connectivity during attention to emotion.

## Methods

### Recruitment

Participants were women aged 18-65 (Mean=42.83, SD=12.38) primarily recruited through the GTP, an ongoing, long-standing collective of trauma and PTSD studies in inner-city Atlanta, Georgia. Participants were approached at random in the waiting rooms of Grady Memorial Hospital medical clinics; recruitment also occurred via flyers distributed in the community, self-referrals through the GTP website, and referrals from clinicaltrials.gov for this study (NCT02754557). Interested individuals underwent informed consent procedures with study staff and subsequently completed a brief battery of questionnaires to assess trauma history and related psychopathology (e.g., PTSD); further details on inclusion/exclusion criteria in Supplement. All procedures were approved by the Emory Institutional Review Board and Grady Research Oversight Committee.

Ninety-three participants were screened and determined to be eligible for the intervention. Five individuals withdrew before starting the intervention, and among the 88 participants that remained, 65 completed the intervention, yielding a 74% retention rate; 31 of these individuals also completed pre- and post-intervention MRI; this is detailed in Figure 1. Some participants started their engagement in the study during the COVID-19 pandemic (n=5). Among the non-completers, one participant withdrew due to safety concerns following the COVID-19 pandemic, one participant withdrew due to receiving a life-threatening health diagnosis, and seven participants withdrew due to lack of continued interest. Four participants were excluded after initial screening: one participant was excluded due to initiating cocaine use and three participants were withdrawn due to prolonged scheduling conflicts. Ten other participants were unresponsive to multiple contact attempts/lost to follow up. Demographic and clinical characteristics of the entire sample are presented in Supplemental Table 1.

**Figure 1.**
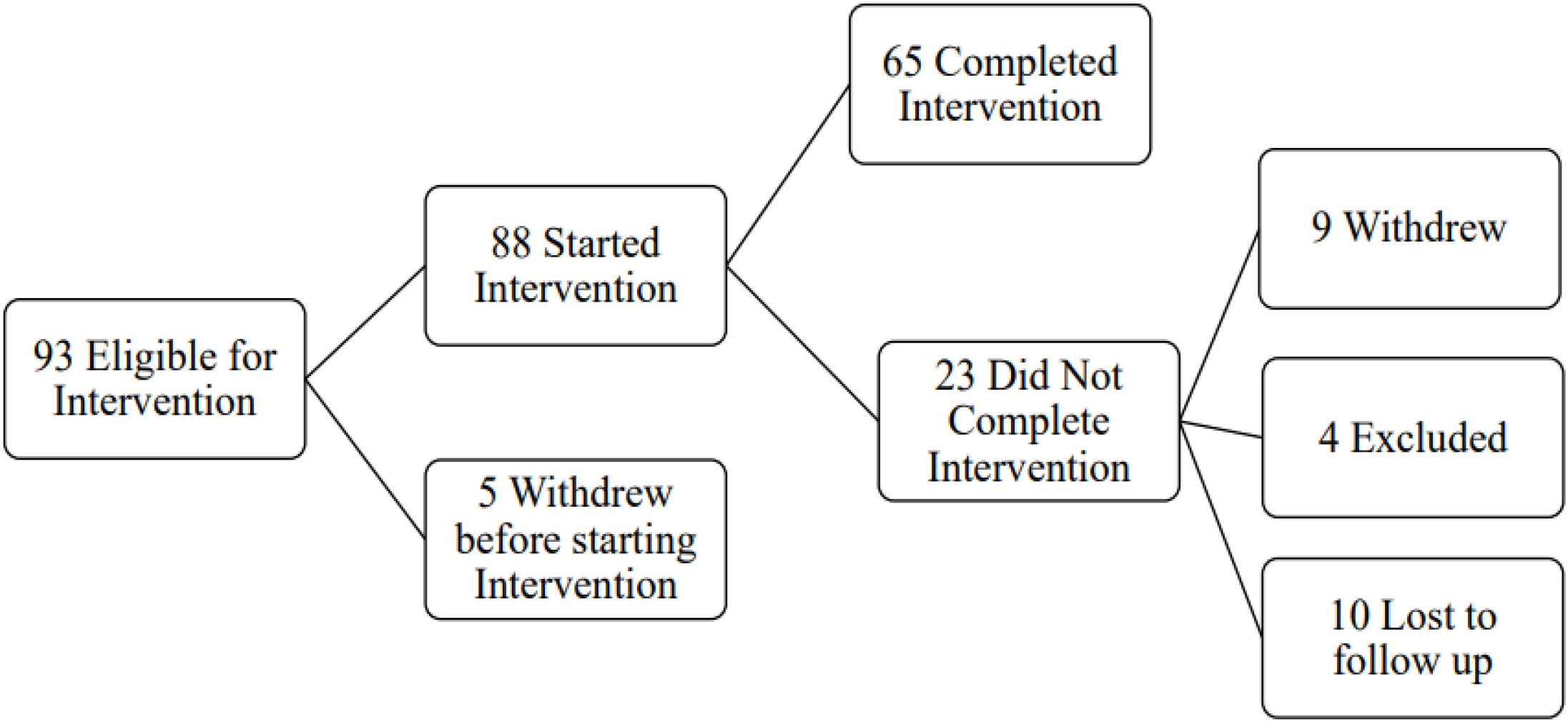
Flowchart of study engagement

Detailed demographic and clinical characteristics of the 65 participants who completed the study are provided in Table 1. A majority of participants (82%) were Black American (1 person identified as Hispanic/Latinx, 7 identified as white, 2 identified as mixed-race and 2 others identified as “other race”) and 65% were significantly economically disadvantaged (monthly household incomes below $2000/month). No significant differences in clinical and demographic characteristics were observed between VBFM and BFM groups at pre-intervention (Table 1 and Supplemental Table 1, *p*s>.05).

**Table 1.** Demographic and Clinical Characteristics

|  | **VBFM** | **BFM** |  |
| --- | --- | --- | --- |
|  | **N=38** | **N= 27** |  |
|  | **% (N)** | **% (N)** | **<u>Fisher's exact</u>** |
| Race |  |  | 1.81 |
| African American/Black | 78.9 (30) | 85.2 (23) |  |
| Hispanic/Latino | 2.6 (1) | 0.0 |  |
| Caucasian/White | 13.2 (5) | 11.1 (3) |  |
| Mixed | 2.6 (1) | 3.7 (1) |  |
| Other | 2.6 (1) | 0.0 |  |
| Household Monthly Income |  |  | 3.42 |
| \$ 0 – 249 | 13.2 (5) | 22.2 (6) | |
| \$ 250 – 499 | 7.9 (3) | 3.7 (1) | |
| \$ 500 – 999 | 23.7 (9) | 18.5 (5) | |
| \$ 1,000 – 1,999 | 13.2 (5) | 25.9 (7) | |
| \$ 2,000 or more | 42.1 (16) | 29.6 (8) | |
| Education |  |  | 4.89 |
| Less than 12<sup>th</sup> grade | 18.4 (7) | 7.4 (2) |  |
| 12<sup>th</sup> grade/ High school graduate | 13.2 (5) | 14.8 (4) |  |
| GED | 10.5 (4) | 3.7 (1) |  |
| Some college or tech school | 10.5 (4) | 18.5 (5) |  |
| Technical school graduate | 7.9 (3) | 14.8 (4) |  |
| College graduate | 26.3 (10) | 18.5 (5) |  |
| Graduate school | 13.2 (5) | 22.2 (6) |  |
| Medication |  |  | **<u>X<sup>2</sup></u>** |
| Antidepressant | 44.7 (17) | 40.7 (11) | .10 |
| Anticonvulsant/Mood Stabilizer | 21.1 (8) | 29.6 (8) | .63 |
| Stimulant | 10.5 (4) | 7.4 (2) | .18 |
| Antipsychotic | 2.6 (1) | 7.4 (2) | .82 |
| Age (Mean, SD, range) | 40.67 (11.91, 19-65) | 45.48 (11.97, 18-63) |  |

### Study Procedures

After the initial screening and consent for this intervention study, participants received a psychodiagnostic interview to rule out exclusionary study diagnoses (e.g., bipolar disorder, schizophrenia); this interview included the MINI neuropsychiatric interview(Sheehan et al., 1998) and Clinician Administered PTSD Scale (CAPS(Weathers et al., 2017), detailed in the Supplement). Data for these measures, including effect sizes (**ꞃp**^**2**^) and correlations of change for all study measures, are provided in Supplemental Tables 2 and 5. After completing the interview, participants were then randomized into one of two BFM intervention conditions using random number assignment. The control group received BFM; the experimental group received VBFM. Electrocardiography (ECG) data were collected during intervention sessions to derive measures of resting HRV

### Intervention Procedures

Full details are provided in Supplement. At each of six intervention visits over ∼three weeks, participants were fitted with ECG leads and sat in a chair in a sound-attenuated chamber in front of a computer screen and microphone. Instructions appeared on the screen, which directed them to either engage in breath focus or rest (1 minute in each condition). The microphone was connected to a Digitech multi-effects board which processed the auditory signal, which was connected to an amplifier and played into a haptic subwoofer that was worn like a pendant on the sternum; the breath waveform was felt as rumbling vibrations that corresponded to the breath. For those receiving the VBFM, breathing into the microphone resulted in a sternal vibration proportional to the person’s breath; participants in the BFM group wore the same device but received no vibratory feedback.

### The Multidimensional Assessment of Interoceptive Awareness

(MAIA(Mehling et al., 2013)) is a 32-item self-report measure of interoceptive awareness with good construct validity and internal consistency(Mehling et al., 2012). The MAIA assesses different facets of interoceptive awareness, including: noticing body signals (Noticing), Not Distracting, Attention Regulation, Emotional Awareness, Self-Regulation, Body-Listening, and Body Trusting. We used an overall summed score as a primary index of interoception; significant between-group findings were subjected to follow-up specificity analyses with MAIA subscales. *Multiscale Dissociation Inventory (MDI)*. The MDI is a 30-item assessment of current dissociative symptoms(Briere, Weathers, & Runtz, 2005); subscales assess different facets of dissociation: emotion disengagement, identity dissociation, memory disturbance, emotional constriction, depersonalization and derealization.

### MRI data acquisition and processing

T1- and T2*-weighted images were acquired on research-dedicated Siemens 3T MRI Systems, detailed in the Supplement.

#### Affective Stroop Functional Connectivity Analyses

The Affective Stroop (AS) task is an attentional control task that has been used in prior studies of PTSD and dissociation(Fani et al., 2019; Fani et al., 2018), detailed in the Supplement. We examined bilateral amygdala connectivity during presentation of all trauma-relevant stimuli and neutral stimuli (including number congruent, number incongruent, and passive view trials) in primary analyses, given our prior research identifying amygdala-insula disconnectivity to trauma-relevant vs neutral stimuli in this measure in dissociative trauma-exposed individuals(Fani et al., 2018). We used a recent comprehensive review of emotion to attention studies to define regions of interest(Dolcos et al., 2020) and conducted seed-to-voxel analyses using the CONN toolbox(Whitfield-Gabrieli & Nieto-Castanon, 2012), detailed in Supplement. At both the pre- and post-intervention timepoints, we examined voxelwise findings at a threshold of p<.05 with family-wise error correction and minimum cluster size of 10mm isotropic. For regions that survived our statistical threshold for both pre- and post-intervention, we extracted average timeseries within these ROIs (left and right) at pre- and post-intervention and entered the values into data analyses.

### Neuropsychological Testing

Forty-two participants were administered two subtests of the Penn Computerized Neuropsychological Battery (CNP)(Gur et al., 2010) to assess sustained attention and working memory: the Continuous Performance Test (CPT) and the Letter N-back (LNB) Task, detailed in the Supplement. Number of correct responses were recorded and analyzed for each task; pre- and post-intervention data in Table 2.

**Table 2.**
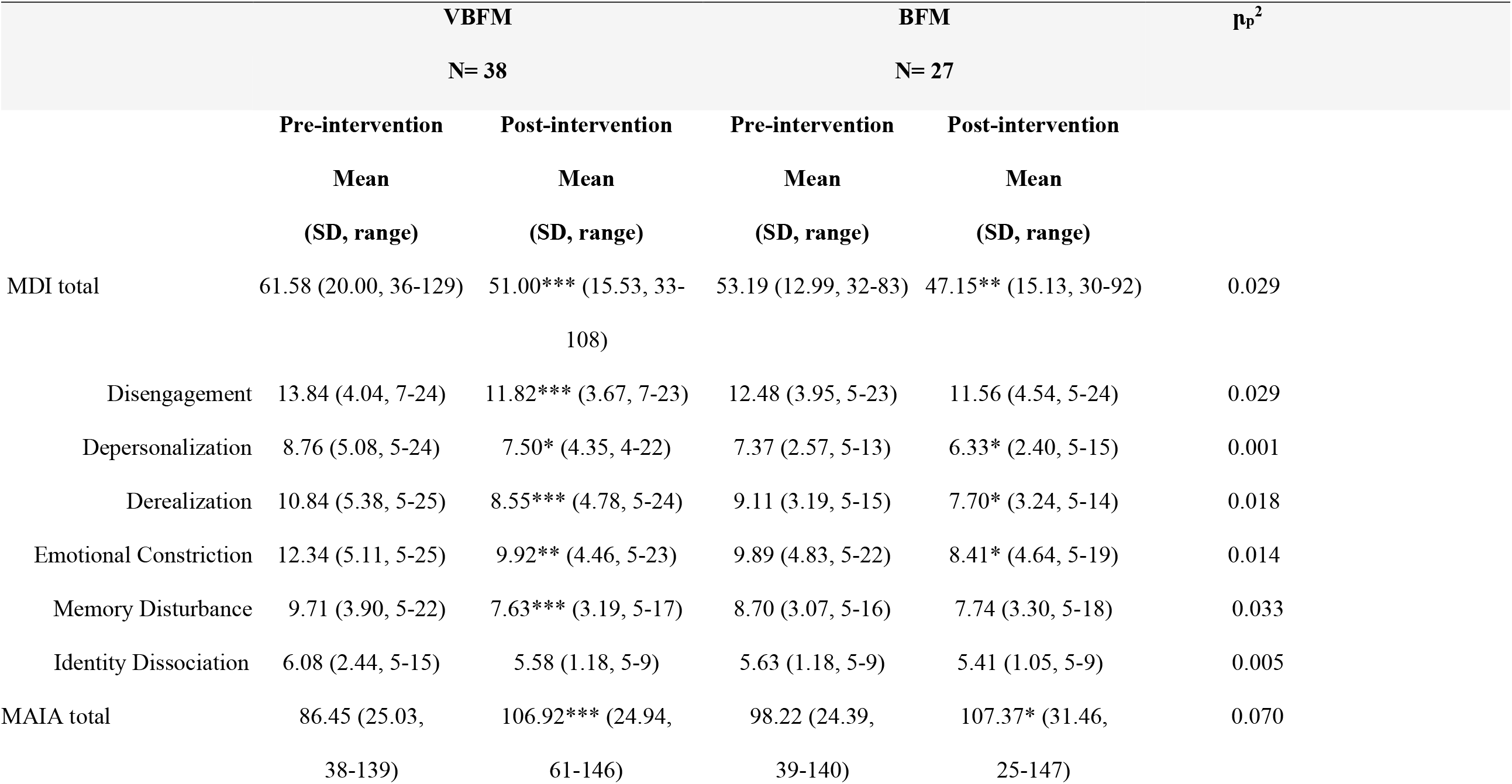
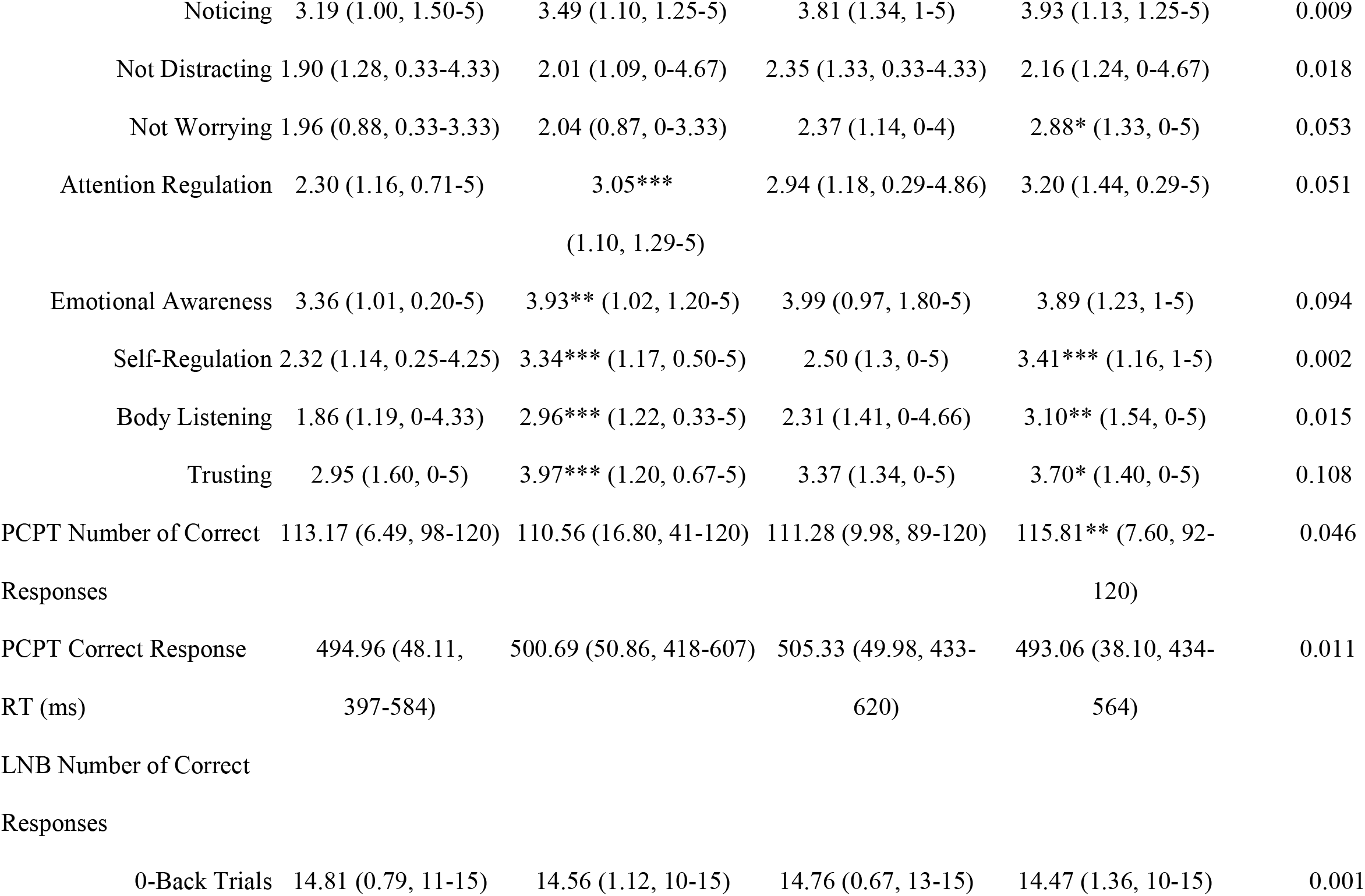
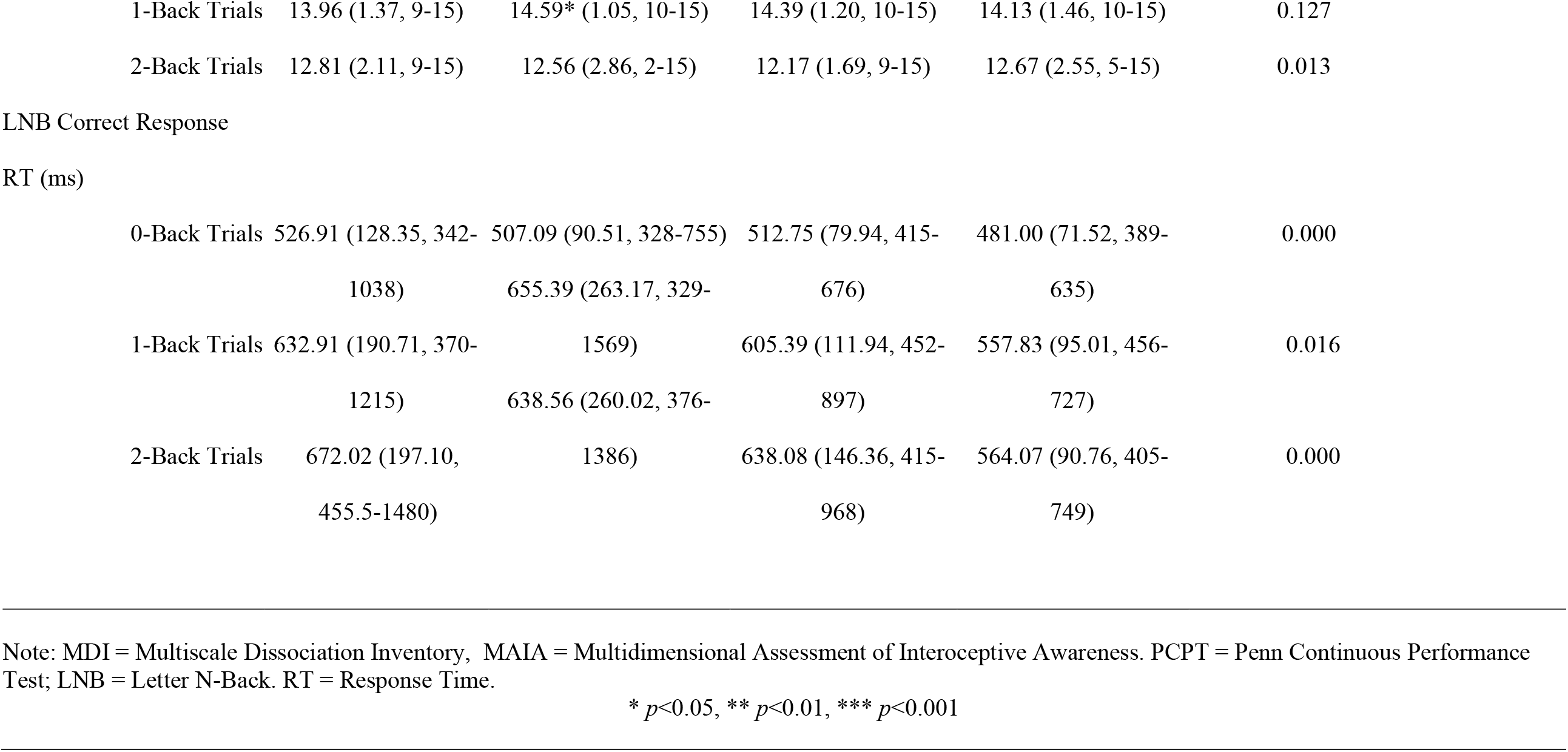
Clinical and Cognitive Assessment Data at Pre- and Post-intervention

### Data Analyses

#### Clinical and neuropsychological data

Three separate mixed model analyses of covariance (ANCOVAs) were used to probe group (VBFM vs BFM) by time (pre-post intervention) interactions with interoceptive awareness (MAIA), attention and cognitive control (CPT, LNB performance). Covariates for all analyses included stimulant medication use, given their direct effects on attention (indicated in 6 participants), study entry during the COVID pandemic (n=5) and race, given the possibility that both the pandemic and race (specifically, racism-related effects) may influence all outcomes of interest, and the fact that racism is a factor that influences attention(Fani, Carter, Harnett, Ressler, & Bradley, 2021; Fani, Jovanovic, et al., 2012; Fani, Tone, et al., 2012); our preference was to examine main and interactive effects of race in our analyses, but the small numbers in the white category (n=7) made meaningful statistical comparisons impossible. Statistical significance for analyses was set at *p*<.05 for each family of tests. <u>AS data</u>. A similar mixed model ANCOVA was conducted to identify group by time by distractor condition interactions for amygdala connectivity from pre-to post-intervention. Similarly, interactions were also probed for AS behavioral responses overall (percent error) for all trials that had a response component (number congruent and number incongruent stimulus trials).

To examine interoception as a potential mechanism of change in dissociation and neurophysiological variables, and to examine whether intervention group moderated these effects, we conducted three separate moderation analyses using mixed model ANCOVAs (same covariates). First, we assessed whether intervention group moderated the relationship between 1) change in interoception (MAIA total change) and change in dissociation (MDI total change); 2) change between interoceptive awareness (MAIA total change) and change in HRV; 3) change in interoception and change in amygdala connectivity. Statistical significance was defined as *p*<.05 for each of these analyses.

## Results

### Interoceptive awareness

No significant effects of time were observed (F_1,60_=.12, p=.728) but a group by time interaction was observed for MAIA total (F_1,60_=4.74, p=.033); the VBFM group demonstrated a greater increase from pre- to post-intervention as compared to BFM (Table 2; Figure 2a). Follow-up analyses with MAIA subscales revealed similar results, with no main effects of time (*p*s>.05) but a group by time interaction for the body trusting (F_1,60_=7.25, p=.009; Figure 2b) and emotional awareness (F_1,60_=6.52, p=.013) subscales.

**Figure 2.**
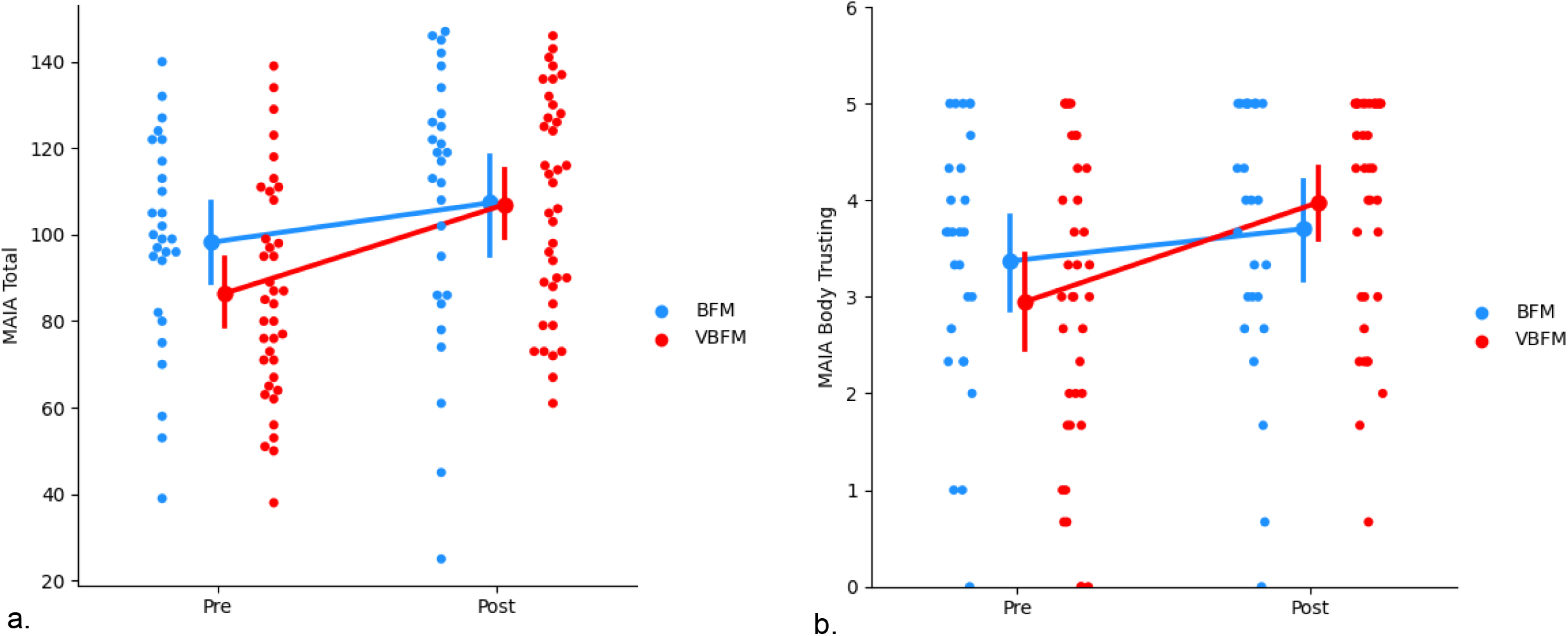
Increases in overall interoception (MAIA total) (a) and body trusting (b) in participants who received vibro-acoustic augmentation during breath focused meditation (VBFM) as compared to participants who engaged in breath focused meditation (BFM) without this augmentation.

### Attention/cognitive Control

#### CPT

No main effects of time (F_1,35_=.12, p=.732) or group by time interaction (F_1,35_=1.63, p=.211) were observed for correct responses on the CPT. Similarly, no main effects of time (F_1,35_=.24, p=.629) or group by time interactions were observed for response time for CPT correct responses (F_1,35_=.01, p=.499). <u>Letter-N-Back</u>. For correct response on the 0-back condition, no main effects of time (F_1,37_=1.19, p=.282) or group by time interactions (F_1,37_=.003, p=.959) were observed. For correct response on the 1-back condition, no main effects of time were observed (F_1,37_=2.70, p=.109), but a group by time interaction was observed (F_1,37_=5.42, p=.025); VBFM participants demonstrated increased correct responses from pre- to post-intervention, whereas BFM participants demonstrated slight decreases in correct responses overall (Table 2).

#### Affective Stroop: Amygdala Connectivity

At pre-intervention at our statistical threshold, significant clusters of increased connectivity emerged in the bilateral temporal lobe, inclusive of hippocampal regions, as well as the left insula; a similar pattern of findings emerged at post intervention, but no insula activation was observed (illustrated in Figure 4a; Supplemental Table 3). No significant clusters emerged in the ACC at either pre or post-intervention or in the bilateral inferior parietal cortex or right insula at post-intervention, which precluded inclusion of these regions in our analyses.

Significant associations between bilateral amygdala connectivity were observed with both the left and right hippocampus at pre- and post-intervention at our statistical threshold (Supplemental Table 3). Repeated measures ANCOVA with left hippocampus connectivity revealed no significant effects of time (F_1,26_=1.48, p=.234) but a significant group by time by condition interaction was observed (F_1,26_=4.83, p=.037). Participants who received the VBFM intervention demonstrated an increase in amygdala-left hippocampus connectivity across both conditions as compared to those with BFM, who demonstrated a decrease in amygdala-left hippocampus connectivity. Significant differences in amygdala-left hippocampus connectivity were observed between the two intervention groups at baseline, with the VBFM group demonstrating overall lower amygdala-left hippocampal connectivity to both trauma-relevant and neutral AS distractor images as compared to the BFM group (F_1,26_=4.99, p=.034). The VBFM group demonstrated an overall increase in amygdala-left hippocampus connectivity whereas the BFM group demonstrated a decrease in connectivity. For amygdala-right hippocampus connectivity, no significant effect of time was observed (F_1,26_=1.51, p=.230) and no group by time interaction was observed (F_1,26_=.04, p=.847).

### Behavioral data

Behavioral data was not recorded for one participant due to a technical failure, leaving n=30 for behavioral data analyses. Repeated measures ANCOVA revealed no significant effects of time (F_1,26_=1.48, p=.234), intervention group by time (F_1,26_=.10, p=.752), or intervention group by time by condition interaction (F_1,26_=.89, p=.356) for percent error on AS trials.

#### Moderation analyses

We tested the hypothesis that the relationship between interoception change and dissociation change would occur as a function of intervention group. A significant interaction of intervention group with change in these variables was observed (F=4.83, p=.032; Figure 3a). Examination of the interaction plot revealed that, for those who received VBFM, increases in interoception corresponded with decreases in dissociation (r=-.47, p=.003), this relationship was not observed in those who did not receive the vibration feedback (r=-.25, p=.207).

**Figure 3.**
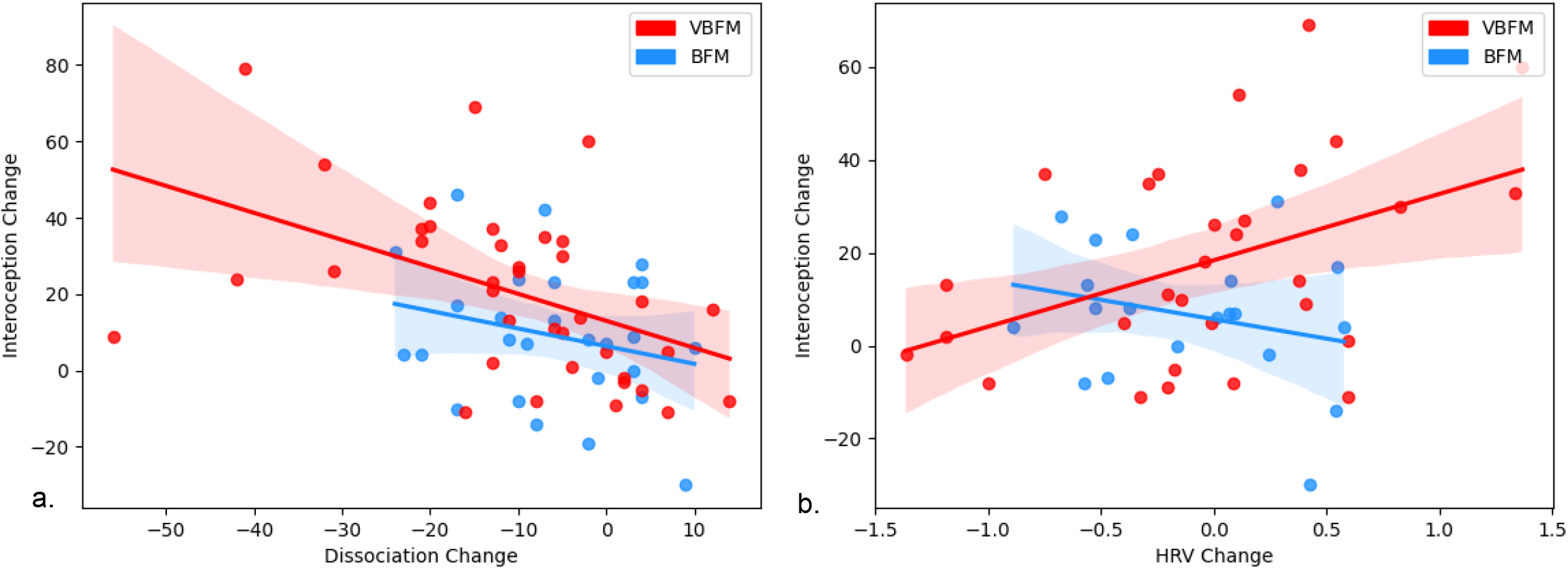
Intervention group moderates the relationship between interoception change and: a) dissociation change; b) high-frequency heart rate variability (HRV) change.

We then tested the hypothesis that the relationship between interoception change and HRV change would occur as a function of intervention group. A significant interaction of intervention group with change in these variables was observed (F=4.27, p=.044; Figure 3b). Examination of the interaction plot revealed that, for those who received VBFM, HRV increased as interoception increased (r=.43, p=.018), a relationship that was not observed among those who received BFM only (r=-.28, p=.27).

Finally, we tested the hypothesis that the relationship between interoception change and amygdala connectivity change (amygdala-left hippocampus connectivity to trauma-relevant or neutral distractor cues) would occur as a function of intervention group. We did not observe a significant interaction of intervention group with change in these variables (F=.53, p=.474). However, bivariate correlations revealed that increased amygdala-left hippocampus connectivity to trauma-relevant distractors was positively associated with interoception change in the VBFM group, (MAIA total; Figure 4b; r=.48, p=.058), particularly body trusting, (r=-.55, p=.027), an association that was not observed in the BFM group (MAIA total r=-.04, p=.91; MAIA body trusting r=-.50, p=.17). No associations between interoception and connectivity were observed for either group in response to neutral AS distractors.

**Figure 4.**
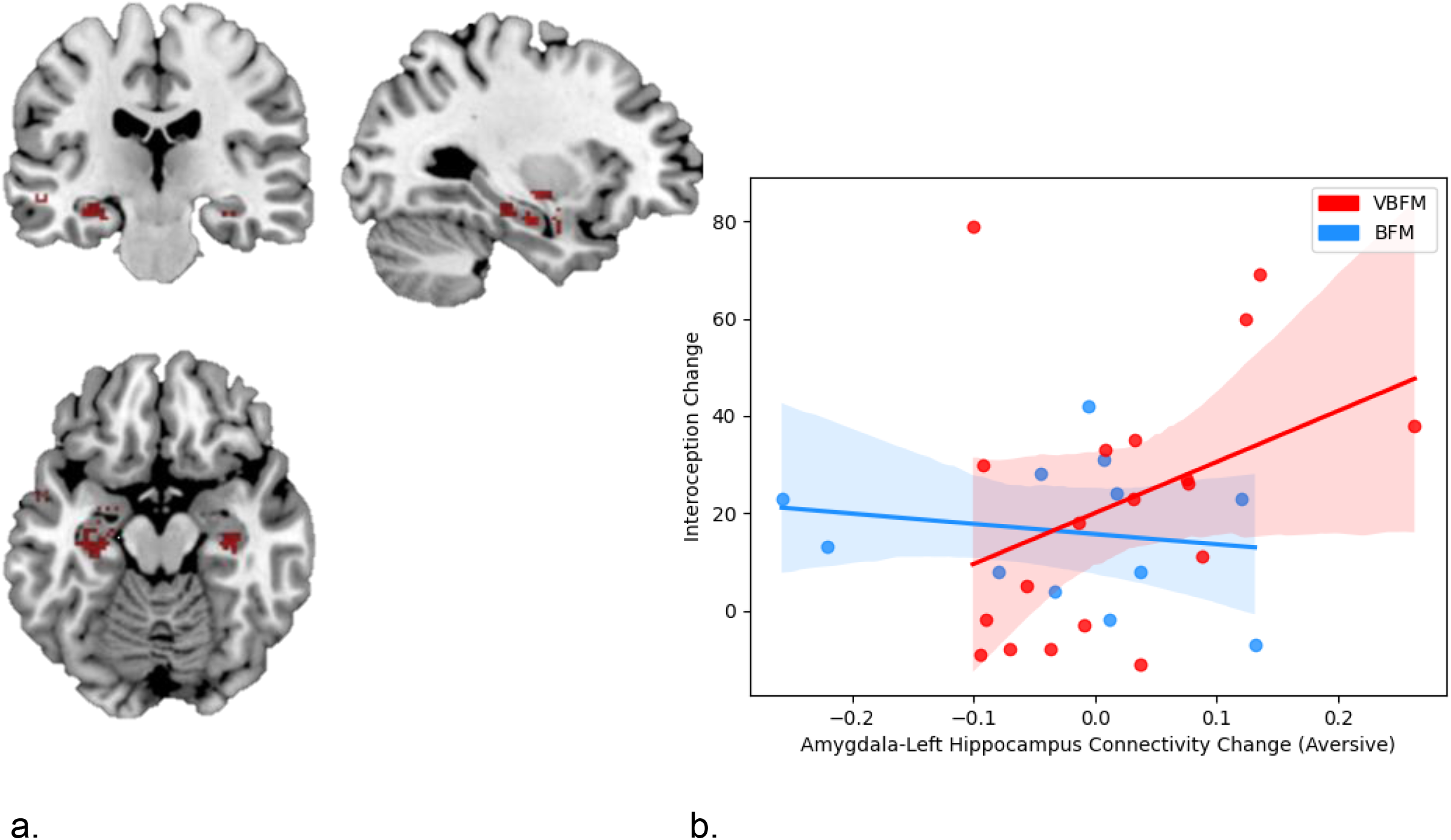
Regions that demonstrated significant connectivity with the amygdala in the entire sample at post-intervention. (b) Increases in amygdala-left hippocampus connectivity to trauma-relevant Affective Stroop distractor stimuli from pre-to post-intervention correspond with increased interoception (MAIA total) in those who received vibro-acoustic augmentation during breath-focused meditation (VBFM), but not in those who engaged in breath focused meditation (BFM) without augmentation.

## Discussion

We tested whether BFM with exteroceptive feedback in the form of vibration proportionate to the breath (VBFM) would elicit changes in interoception and attentional control, autonomic regulation (HRV) and amygdala connectivity in a sample of trauma-exposed women with dissociative symptoms, a majority of whom were Black. We also examined whether intervention condition would moderate the relationship between changes in dissociation, HRV, and amygdala connectivity. We found that participants who received vibration feedback during BFM demonstrated greater improvements in interoception, particularly body trusting and emotional awareness, and sustained attention on a neuropsychological measure. They also demonstrated increased amygdala-left hippocampus connectivity. In the VBFM group only, HRV increased as interoception increased, and interoception changes corresponded with greater improvements in dissociation. These data provide evidence for interoception being a mechanism of change with VBFM, and that this mechanism translated to neurophysiological changes.

To our knowledge, our study is the first to demonstrate the effects of exteroceptive augmentation of a mindfulness intervention on interoception, attention, autonomic regulation and neural connectivity in a trauma-exposed population with dissociative symptoms. Exteroceptive augmentation has been shown to improve interoception and concurrent physiology(Suzuki et al., 2013), and MBIs improve interoception(Farb et al., 2013; Kang, Sponheim, & Lim, 2022; Lima-Araujo et al., 2022). Some trauma therapies involve drawing the patient’s attention to internal visceral or musculo-skeletal sensations (e.g., Somatic Experiencing therapy(Payne, Levine, & Crane-Godreau, 2015)). It appears that exteroceptive augmentation of these signals can facilitate attentional focus and enhance interoceptive functions.

Participants who received VBFM demonstrated particularly greater increases in the body trusting aspect of interoception—i.e., the ability to experience the body as trustworthy and safe. Difficulties with body trusting are a common consequence of trauma, particularly trauma that has an interpersonal component(Vanderkolk, 1994); these difficulties are not limited to dissociation(Dunne, Flores, Gawande, & Schuman-Olivier, 2021; Mehling et al., 2013). Notably, associations between body trusting difficulties and depression have also been shown in Black women with depression and hypertension(Solano Lopez & Moore, 2019). Impairments in body trusting may preclude other aspects of interoception, including body awareness and interoceptive prediction, and lead to dysregulation in emotion and homeostasis/autonomic regulation. Given such findings(Solano Lopez & Moore, 2019) these problems may be even more likely to disrupt autonomic regulation in Black women, who are already disproportionately burdened with trauma(Gillespie et al., 2009; Gluck et al., 2021) and psychosocial stressors such as gendered racism and structural inequities(Lee, Perez, Boykin, & Mendoza-Denton, 2019). As such, VBFM could provide a unique way to enhance the ability to trust body signals, which, in turn, may augment functioning of regulatory homeostatic mechanisms.

In support of this notion, we found that improvements in autonomic regulation were associated with improved interoception only in VBFM participants. The addition of a sensory stimulus—vibration—to a self-regulatory practice, BFM, may be a pathway toward improving autonomic regulation via interoceptive enhancement. Consistent with polyvagal theory, which indicates that autonomic regulation is intertwined with emotional states and social behaviors(Porges, 2007, 2009), our findings suggest that improvements in interoception produced by exteroceptively-augmented self-regulation practice supports improved autonomic nervous system regulation, and flexible, adaptive responding to stressors. These changes, in turn, may promote various facets of emotional and physical well-being that encompass physiology, behavior, and clinical symptoms. Our finding that interoceptive improvements corresponded with proportionately greater improvements in dissociation in VBFM vs BFM participants supports this idea.

As predicted, participants who received VBFM also demonstrated greater improvements in sustained attention on a neuropsychological (N-back) task. Vibration itself appears to have an impact on attentional processes; whole body vibration has been shown to increase attentional vigilance(Poulton, 1978) and improved Stroop task performance in attention deficit hyperactivity disorder(Fuermaier et al., 2014); forearm vibration has produced improved performance on a target detection task and shorter reaction times in patients with traumatic brain injury(Muller et al., 2002). It appears that exteroceptive feedback in the form of vibration during a type of attention training, breath-focused mindfulness, may be a way to augment sustained attention in individuals with attention dysregulation, including dissociative individuals. Attentional orienting is easily guided by sensory stimulation; therapies such as eye movement desensitization and reprocessing capitalize on this, employing side-to-side movements of light or other visual stimuli during therapeutic exposures that involve recollection of traumatic events. When coupled with an interoceptively-focused self-regulation practice, vibration may facilitate attentional focus, addressing a major barrier to treatment engagement in symptomatic trauma-exposed individuals.

VBFM participants also demonstrated an increase in connectivity between the amygdala and left hippocampus, key emotion regulation and interoceptive network nodes, during AS performance. Changes in connectivity corresponded with changes in interoception for only participants who received VBFM, showing further evidence that interoception is a primary change mechanism. Evidence from animal models(Clifton, Vickers, & Somerville, 1998; Davidson & Jarrard, 1993; Lathe, Singadia, Jordan, & Riedel, 2020) and lesion studies indicate that the hippocampus is essential to interoceptive functions; the famous patient H.M. who received bilateral temporal lobe resection for epilepsy was not able to detect bodily states, including pain, hunger and thirst(Hebben, Corkin, Eichenbaum, & Shedlack, 1985). Further, there is evidence for the hippocampus’ role in integrating exteroceptive and interoceptive information in the formation of memories(Kassab & Alexandre, 2015); the hippocampi bind together perceptual aspects of experience with emotional valence features, which is driven by reciprocal connections to the amygdala(Pitkanen, Pikkarainen, Nurminen, & Ylinen, 2000). As such, increased connectivity between the amygdala and hippocampus in those who received VBFM could indicate improved integration of attention and interoceptive processing in these individuals.

Further, given that our participants were mostly Black individuals, our findings add to a growing literature indicating the utility of mindfulness-based interventions in Black individuals and the possibility that this intervention strategy may be useful for individuals from marginalized racial and ethnic groups(Dawson, Jones, Fairbairn, & Laurent, 2022). As such, these data show evidence that vibroacoustic enhancement of breath-focused mindfulness may be an efficient and culturally-acceptable intervention for people from racially marginalized groups, although qualitative data is needed to make conclusive statements on acceptability.

We acknowledge some study limitations. First, our sample size for psychophysiology, neuropsychological and neuroimaging data was relatively limited, which consequently limited our statistical power for moderation analyses. This could be an explanation for non-significant differences in performance on the AS, as well as non-significant pre-post-intervention changes in amygdala connectivity to the insula, a major hub for interoceptive processes. It should be noted that this was a pilot study with limited resources for MRIs; nonetheless, the direction of findings was largely consistent with our hypotheses, and we are currently conducting a larger study across different sites to comprehensively examine neurophysiological mechanisms of VBFM (NCT04670640). Finally, the fact that we included all women and majority Black Americans may be perceived as a limitation in generalizability; however, we believe this to be a strength, given the significant lack of representation of Black women in mechanistic clinical trials.

In conclusion, we found that that breath-synced vibration produced enhanced interoception, and these enhancements appeared to affect autonomic regulation, attentional control, symptoms of dissociation, and increased connectivity in neural systems involved with interoception and emotion regulation. These promising, converging lines of data indicate that a relatively simple augmentation of a common mindfulness intervention—BFM—may help dissociative individuals overcome trauma-focused treatment barriers. This straightforward technology could be used to produce a commercially-available device and application that can be used on demand, to assist with common trauma-related problems such as dissociation and attentional control difficulties. The data presented here point to new directions in device-assisted mental health interventions that can be made accessible to those who are most in need, particularly those with limited mental health care access.

## Funding and Acknowledgments

This work was supported by National Institutes of Mental Health (MH101380 to NF, HD071982 to BB), the National Center for Complementary and Integrative Health (AT011267 to NF and GJS), the Emory Medical Care Foundation and Emory University Research Council, the American Psychological Association, Society for Clinical Neuropsychology and the Georgia Institute of Technology/Georgia State University Center for Advanced Brain Imaging. We wish to thank Allen Graham, Rebecca Hinrichs, Angelo Brown and other members of the Grady Trauma Project, as well as members of the Fani laboratory for their assistance with data collection. We thank all of our participants for their time and involvement in this study.

