## Supplement for "Neurophysiological Changes Associated with Vibroacoustically-augmented Breath-focused Mindfulness for Dissociation: Targeting Interoception and Attention"

Supplemental Methods

*Recruitment*. Exclusion criteria for GTP studies were as follows: 1) younger than 18 years of age or older than 65 years of age; 2) currently at risk for suicide, experiencing active psychosis, or cognitively compromised (e.g., intoxication, severe intellectual disability). All participants were screened over the phone to assess eligibility for this study. Since participants were largely recruited from GTP and extant projects were recruiting women only (e.g., MH101380), only women were recruited for the present study. Participants were included in this study based on the following criteria: 1) experienced at least one PTSD Criterion A traumatic stressor, according to the DSM-IV or DSM-5; 2) presence of current, impairing trauma-related PTSD symptoms (defined as meeting diagnostic criteria for at least two of the four PTSD symptom clusters), and 3) willingness to participate in the study. Exclusion criteria for the intervention study were the following: 1) a history of bipolar disorder, schizophrenia or primary psychotic disorder; 2) a history of neurological conditions or injuries including strokes, seizures, and traumatic brain injury, or history of any disorder with central nervous system involvement (e.g., HIV); 3) current substance or alcohol dependence.

*Intervention Procedures*. At each of six intervention visits over approximately three weeks, participants were fitted with electrophysiology leads and sat in a chair in a sound-attenuated chamber in front of a computer screen and microphone while being monitored by a researcher. Instructions appeared on the screen, which instructed them to either engage in breath focus or rest, for 1 minute in each condition. These conditions were randomized; the total session time was 15 minutes. The microphone was connected to a Digitech multi-effects board which processed the auditory signal (amplified it, smoothed transients, sustain detected signals, and dropped the ensuing waveform by an octave), which was connected to an amplifier and played into a haptic subwoofer (Dayton Audio TT25-8 Puck) that was worn like a pendant on the participant’s sternum; the effect was that the breath waveform was felt as rumbling vibrations that corresponded to the breath. Study staff ensured that the participant understood the instructions fully prior to starting data collection. For those receiving the VBFM, breathing into the microphone resulted in a vibration on their sternum that was proportional to the person’s breath; individuals who were in the BFM group wore the same device but received no vibratory feedback.

*Electrocardiography (ECG) Data collection.* ECG data were collected during both rest and active intervention (BFM or VBFM) conditions to derive measures of HRV; resting HRV data were analyzed here. ECG data were derived from a two-lead montage (right neck, left wrist; 2000 Hz, no filters), using a BIOPAC ECG100C Electrocardiogram amplifier (BIOPAC Systems, Inc) via AcqKnowledge software version 4.0. *Processing.* ECG data were exported to text files which were processed and cleaned using via custom Matlab code. R-wave peaks were extracted and converted to inter-beat interval series for each block. Series were subjected to continuous Morelet waveform transforms to yield a block-wise running estimate of power in the high frequency heart rate variability (HF-HRV) band (0.18-0.4 Hz) using the HRV-AS package^1^, available at <https://github.com/jramshur/HRVAS>). Quality assurance metrics included: heart rate between 40 and 120 beats per minute; no significant abrupt spikes or ectopic beats; electrocardiograms that appear normative with distinct R peaks and QRS complexes. Data that fell outside expected values were manually checked and corrected. Pre- and post-intervention HRV data for 50 participants survived quality assurance checks. Mean HF HRV was calculated at rest for each participant; pre- and post intervention resting HRV was Winsorized before being entered into group analyses.

*Mini-International Neuropsychiatric Interview for DSM-IV* (MINI). The MINI international neuropsychiatric interview version 6^2^ is a psychometrically sound, structured clinician-administered interview which was used to assess for lifetime substance use history presence and severity; presence of mood disorder (bipolar and unipolar depression), lifetime PTSD, and psychotic disorders. *The Clinician Administered PTSD Scale-version 5 (CAPS-5)*^3^ is a structured, clinician-administered interview that was used to assess for the presence and severity of PTSD, considered a gold standard PTSD assessment with excellent psychometric properties^3^. All interviews were performed by study staff under the supervision of licensed psychologists (NF, AP). CAPS-5 and MINI data for the sample are provided in Supplemental Table 2.

*Affective Stroop Task*. The Affective Stroop (AS) task comprises 16 trials for each of three conditions: number congruent, number incongruent, and passive viewing of each of three types of distractor images, trauma-relevant, positive, and neutral, as shown in Supplemental Figure 1. A total of 40 fixation trials (2500 ms duration) are also randomly presented for a total of 144 trials. Distractor and number stimuli are each presented for 400 ms, followed by a blank screen (1300 ms). Each trial includes two identical distractor stimuli. Distractor images were unique for each trial condition. The task includes images relevant to traumatic experiences of our study population, based on our earlier studies, which indicate a high amount of interpersonal trauma and assault involving a weapon ^4^. Trauma-relevant stimuli included scenes of assault and gun violence—all but one of these scenes included people in the images. Approximately half of the positive stimuli were pictures that included people (43%), a minority included animals (17%), activities (theme park, 6%; sports, 23%) or nature scenes (12%). Neutral stimuli included a similar proportion of person-related images (36%) and nature scenes (14%) but also included object stimuli (50%). A majority of the images were obtained from the International Affective Picture System IAPS; ^5^. Practice trials (with a separate set of distractor images) were administered to all participants prior to the scan, and task proficiency (accuracy on a majority of trials) was required before actual task administration. First-level models were created for each participant for each emotion (trauma-relevant, positive, neutral) and cognitive condition (number incongruent, number congruent, passive viewing); we examined amygdala connectivity during all trauma-relevant stimuli and neutral stimuli in primary analyses, given our prior research identifying amygdala-insula dysconnectivity to aversive vs neutral stimuli in this measure in dissociative trauma-exposed individuals ^6^.

The Penn Computerized Neuropsychological Battery (CNP) was administered to 42 participants; this battery has demonstrated good reliability and construct validity in psychiatric and non-psychiatric populations^7^

*The Penn Continuous Performance Test (PCPT)^8^* assays sustained attention, measuring errors of omission (inattention) and commission (impulsivity) similar to other continuous performance tests, such as Conners’ Continuous Performance Test II ^9^. Participants are presented with two sets of stimuli: a series of numbers and non-number distractors; a series of letters and non-letter distractors, each presented at 1 s duration). They are asked to press a button when they see an actual number or letter. Participants must successfully complete a brief practice round before moving on to the actual task. Number of correct responses were recorded and analyzed for each of the three conditions. The PCPT demonstrates a moderate correlation with the Penn Letter N-back task (CPT number condition, r= .5, CPT letter condition, R= .6), according to a validation study ^7^. Data from two participants was incomplete due to technical issues, leaving a final sample of n=40 for this measure.

*Penn Letter N-Back Task (LNB).* This task assesses attention and working memory. A series of letters are presented at 2.5 second durations; participants are asked to press a button to indicate their response under three different rules: the 0-back condition, which requires participants to press a button when they see the letter “X”; the 1-back and 2-back conditions, which requires participants to press a button when they view a letter that appeared 1-back or 2-back from the current letter, respectively. A practice round for each condition must be successfully completed before progressing to the actual task. Number of correct responses were recorded and analyzed for each condition.

*MRI data acquisition*

Scans were acquired on three Siemens 3T MRI systems. A list of parameters and number of subjects scanned on each system is listed in Supplemental Table 4. Each subject received their pre- and post-intervention scans on the same scanner.

*MRI processing*

A high-resolution T1-weighted structural scan was acquired for co-registration purposes using an MPRAGE sequence (176 slices; 1 mm isotropic voxels; repetition time (TR) = 2600 msec; echo time (TE) = 3.02 msec; inversion time (TI) = 900 msec; flip angle = 8°). Functional images (190 volumes) were acquired during task administration using a T2*-weighted gradient echo planar sequence (40 interleaved transaxial slices; 3 mm isotropic voxels; TR= 2500 msec; TE = 30 msec; flip angle = 90°). Scans were acquired on three different Siemens 3T MRI systems (n=5/8/18). Sequence parameters were altered slightly to accommodate different models of each scanner (see Supplement for details). A standardized preprocessing pipeline for fMRI data, fMRIprep v20.2.3 (https://fmriprep.org) was used for image pre-processing^10^ based on Nipype 1.6.1^11^. Pre-processing steps involved slice-time and motion correction, co-registration with the T1-weighted scan, normalization to standard space (International Consortium for Brain Mapping 152 Nonlinear Asymmetrical template version 2009c), and segmentation, further detailed below.

*Anatomical data preprocessing*

One T1-weighted (T1w) images was found for each participant within the input BIDS dataset. The T1-weighted (T1w) image was corrected for intensity non-uniformity (INU) with N4BiasFieldCorrection (Tustison et al. 2010), distributed with ANTs 2.3.3 (Avants et al. 2008, RRID:SCR_004757), and used as T1w-reference throughout the workflow. The T1w-reference was then skull-stripped with a Nipype implementation of the antsBrainExtraction.sh workflow (from ANTs), using OASIS30ANTs as target template. Brain tissue segmentation of cerebrospinal fluid (CSF), white-matter (WM) and gray-matter (GM) was performed on the brain-extracted T1w using fast (FSL 5.0.9, RRID:SCR_002823, Zhang, Brady, and Smith 2001). Brain surfaces were reconstructed using recon-all (FreeSurfer 6.0.1, RRID:SCR_001847, Dale, Fischl, and Sereno 1999), and the brain mask estimated previously was refined with a custom variation of the method to reconcile ANTs-derived and FreeSurfer-derived segmentations of the cortical gray-matter of Mindboggle (RRID:SCR_002438, Klein et al. 2017). Volume-based spatial normalization to two standard spaces (MNI152NLin2009cAsym, MNI152NLin6Asym) was performed through nonlinear registration with antsRegistration (ANTs 2.3.3), using brain-extracted versions of both T1w reference and the T1w template. The following templates were selected for spatial normalization: ICBM 152 Nonlinear Asymmetrical template version 2009c [Fonov et al. (2009), RRID:SCR_008796; TemplateFlow ID: MNI152NLin2009cAsym], FSL\u2019s MNI ICBM 152 non-linear 6th Generation Asymmetric Average Brain Stereotaxic Registration Model [Evans et al. (2012), RRID:SCR_002823; TemplateFlow ID: MNI152NLin6Asym].

*Functional data preprocessing*

For each BOLD run per subject (across all tasks and sessions), the following preprocessing was performed. First, a reference volume and its skull-stripped version were generated using a custom methodology of fMRIPrep. Susceptibility distortion correction (SDC) was omitted. The BOLD reference was then co-registered to the T1w reference using bbregister (FreeSurfer) which implements boundary-based registration (Greve and Fischl 2009). Co-registration was configured with six degrees of freedom. Head-motion parameters with respect to the BOLD reference (transformation matrices, and six corresponding rotation and translation parameters) are estimated before any spatiotemporal filtering using mcflirt (FSL 5.0.9, Jenkinson et al. 2002). BOLD runs were slice-time corrected using 3dTshift from AFNI 20160207 (Cox and Hyde 1997, RRID:SCR_005927). The BOLD time-series (including slice-timing correction when applied) were resampled onto their original, native space by applying the transforms to correct for head-motion. These resampled BOLD time-series will be referred to as preprocessed BOLD in original space, or just preprocessed BOLD. The BOLD time-series were resampled into standard space, generating a preprocessed BOLD run in MNI152NLin2009cAsym space. First, a reference volume and its skull-stripped version were generated using a custom methodology of fMRIPrep. Automatic removal of motion artifacts using independent component analysis (ICA-AROMA, Pruim et al. 2015) was performed on the preprocessed BOLD on MNI space time-series after removal of non-steady state volumes and spatial smoothing with an isotropic, Gaussian kernel of 6mm FWHM (full-width half-maximum). Corresponding \u201cnon-aggresively\u201d denoised runs were produced after such smoothing. Additionally, the \u201caggressive\u201d noise-regressors were collected and placed in the corresponding confounds file. Several confounding time-series were calculated based on the preprocessed BOLD: framewise displacement (FD), DVARS and three region-wise global signals. FD was computed using two formulations following Power (absolute sum of relative motions, Power et al. (2014)) and Jenkinson (relative root mean square displacement between affines, Jenkinson et al. (2002)). FD and DVARS are calculated for each functional run, both using their implementations in Nipype (following the definitions by Power et al. 2014). The three global signals are extracted within the CSF, the WM, and the whole-brain masks. Additionally, a set of physiological regressors were extracted to allow for component-based noise correction (CompCor, Behzadi et al. 2007). Principal components are estimated after high-pass filtering the preprocessed BOLD time-series (using a discrete cosine filter with 128s cut-off) for the two CompCor variants: temporal (tCompCor) and anatomical (aCompCor). tCompCor components are then calculated from the top 2% variable voxels within the brain mask. For aCompCor, three probabilistic masks (CSF, WM and combined CSF+WM) are generated in anatomical space. The implementation differs from that of Behzadi et al. in that instead of eroding the masks by 2 pixels on BOLD space, the aCompCor masks are subtracted a mask of pixels that likely contain a volume fraction of GM. This mask is obtained by dilating a GM mask extracted from the FreeSurfer\u2019s aseg segmentation, and it ensures components are not extracted from voxels containing a minimal fraction of GM. Finally, these masks are resampled into BOLD space and binarized by thresholding at 0.99 (as in the original implementation). Components are also calculated separately within the WM and CSF masks. For each CompCor decomposition, the k components with the largest singular values are retained, such that the retained components\u2019 time series are sufficient to explain 50 percent of variance across the nuisance mask (CSF, WM, combined, or temporal). The remaining components are dropped from consideration. The head-motion estimates calculated in the correction step were also placed within the corresponding confounds file. The confound time series derived from head motion estimates and global signals were expanded with the inclusion of temporal derivatives and quadratic terms for each (Satterthwaite et al. 2013). Frames that exceeded a threshold of 0.5 mm FD or 1.5 standardised DVARS were annotated as motion outliers. All resamplings can be performed with a single interpolation step by composing all the pertinent transformations (i.e. head-motion transform matrices, susceptibility distortion correction when available, and co-registrations to anatomical and output spaces). Gridded (volumetric) resamplings were performed using antsApplyTransforms (ANTs), configured with Lanczos interpolation to minimize the smoothing effects of other kernels (Lanczos 1964). Non-gridded (surface) resamplings were performed using mri_vol2surf (FreeSurfer).

Many internal operations of fMRIPrep use Nilearn 0.6.2 (Abraham et al. 2014, RRID:SCR_001362), mostly within the functional processing workflow. For more details of the pipeline, see the section corresponding to workflows in fMRIPrep\u2019s documentation.

*Affective Stroop Seed-based Functional Connectivity Analyses*. We used a recent comprehensive review of emotion to attention studies to define regions of interest for this study^12^; as the amygdala was identified as a critical hub for emotion-related processing, we used the bilateral amygdala as our seed for connectivity analyses, and conducted seed-to-voxel analyses with the CONN toolbox^13^. At both the pre- and post-intervention timepoints, we examined voxelwise findings at a threshold of p<.05 with family-wise error (FWE) correction and minimum cluster size of 10mm isotropic. We identified the insula, inferior parietal cortex, bilateral hippocampus, and anterior cingulate cortex (ACC) as regions of interest (ROIs) given their identified relevance for attentional control in the context of emotion, as well as mindfulness practices^12, 14^; we used masks created from the Wake Forest University PickAtlas to define our ROIs^15^. For regions that survived our statistical threshold for both pre- and post-intervention, we extracted average timeseries within these ROIs (left and right, examined separately) at pre- and post-intervention and entered the values into a mixed model ANCOVA to identify group by time by condition interactions for amygdala connectivity from pre- to post-intervention; covariates are identified in the *Data Analyses* section. Significant interactions were subject to two follow up moderation analyses to examine whether intervention group moderated changes in 1) interoception or 2) dissociation.

*Seed-based Functional Connectivity Analyses*

The functional connectivity analyses were carried out using the CONN Toolbox v21b ^13^. The output images from fMRIPrep (non-ICA-AROMA denoised) were imported into CONN, smoothed with a 6mm Gaussian kernel and denoised. The resulting images were entered into a seed-based connectivity model using the bilateral amygdala as a seed. The amygdala ROI was generated from the CIT168 probabilistic atlas (Tyszka and Pauli, 2016).

**Results**

Interoceptive awareness. For the MAIA not worrying subscale there was no main effect of time (F_1,60_=1.25, p=.27) but a marginally significant group by time interaction was observed (F_1,60_=3.79, p=.056). A similar non-significant group by time interaction was observed for the MAIA attention regulation subscale (F_1,60_=3.33, p=.073), with no significant main effects of time. There were no significant main effects of time or group by time interactions for the other MAIA subscales (*p*s>.05).

*Attention/cognitive Control*: CPT. Given that data showed evidence of non-normal distribution at pre-intervention (Shapiro Wilk statistic=.82, p<.001) and post-intervention (Shapiro Wilk statistic=.54, p<.001) due to the presence of three outliers, data analyses were repeated without outliers. Again, no main effects of time (F_1,32_=.61, p=.44) or group by time interactions (F_1,32_=1.32, p=.260) were observed for CPT correct responses; similarly, no main effects of time (F_1,32_=.24, p=.627) or group by time interactions (F_1,32_=.26, p=.615) were observed for response time on CPT correct responses.

Letter-N-Back. Given the presence of three outliers in the pre- and post-intervention responses and its effects on normal distribution of pre (Shapiro Wilk statistic=.65, p<.001) and post-intervention responses (Shapiro Wilk statistic=.74, p<.001) we repeated analyses without these outliers, and findings were slightly stronger (group by time F_1,34_=7.35, p=.010). For correct response on the 2-back condition, no main effects of time (F_1,37_=1.12, p=.298) or group by time interactions (F_1,37_=.97, p=.332) were observed.

*Dissociation*. Although mean scores were somewhat lower from pre- to post-intervention, no main effect of time (F_1,60_=2.62, p=.11) or group by time interaction was observed for MDI total score (F_1,60_=1.87, p=.18). MDI subscales showed similar decreases from pre- to post-intervention that were not statistically significant (all *p*s>.05). Given non-normal pre-intervention distribution of MDI total score (Shapiro Wilk statistic=.876, p<.001), these scores were natural log transformed and analyses repeated; findings remained similar, with both groups showing a non-statistically significant decrease in overall dissociation severity from pre- to post-intervention (F_1,60_=3.69, p=.060) and no significant group by time interaction (F_1,60_=.73, p=.395).

| *Supplemental. Table 1. Demographic Characteristics: Intent to Treat Comparisons* | | | |
| --- | --- | --- | --- |
|  | **Discontinued Study**  **N=23** | **Completed Study**  **N=65** | **Fishers Exact Test** |
|  | **% (N)** | **% (N)** |  |
| Race  African American/Black  Asian  Hispanic/Latino  Caucasian/White  Mixed  Other | 69.6 (16)  4.3 (1)  0.0 (0)  17.4 (4)  4.3 (1)  4.3 (1) | 81.5 (53)  0.0 (0)  1.5 (1)  12.3 (8)  3.1 (2)  1.5 (1) | 5.06 |
| Household Monthly Income^1^  $ 0 – 249  $ 250 – 499  $ 500 – 999  $ 1,000 – 1,999  $ 2,000 or more | 14.3 (3)  4.8 (1)  0.0 (0)  38.1 (8)  42.9 (9) | 16.9 (11)  6.2 (4)  21.5 (14)  18.5 (12)  36.9 (24) | 8.04 |
| Education^2^  Less than 12^th^ grade  12^th^ grade/ High school graduate | 4.5 (1)  9.1 (2) | 13.8 (9)  13.8 (9) | 5.988 |
| GED  Some college or technical school  Technical school graduate  College graduate  Graduate school | 9.1 (2)  36.4 (8)  9.1 (2)  13.6 (3)  18.2 (4) | 7.7 (5)  13.8 (9)  10.8 (7)  23.1 (15)  16.9 (11) |  |
|  | **Mean (SD, range)** | |  |
| Age | 38.00 (12.10, 19-57) | 42.52 (12.28, 18-25) |  |
| ^1^ Missing data on two participants in the Discontinued group | | | |
| ^2^ Missing data on one participant in the Discontinued group | | | |

| *Supplemental Table 2. MINI and CAPS data* | | |  |
| --- | --- | --- | --- |
|  | **VBFM**  **N=38** | **BFM**  **N= 27** |  |
|  | **% (N)** | **% (N)** | **Pearson Chi-Square** |
| Current Major Depressive Disorder  Lifetime Major Depressive Disorder  Current Substance Use Disorder^1^  Lifetime Substance Use Disorder^1^ | 57.9 (22)  81.6 (31)  21.6 (8)  45.9 (17) | 44.4 (12)  66.7 (18)  0.0  22.2 (6) | 1.15  1.89  6.67  3.82 |
| Current Alcohol Use Disorder^1^  Lifetime Alcohol Use Disorder^1^ | 8.1 (3)  35.1 (13) | 0.0  18.5 (5) | 2.30  2.13 |
| PTSD Symptom Severity Total  Cluster B Symptom Severity  Cluster C Symptom Severity  Cluster D Symptom Severity  Cluster E Symptom Severity  Cluster G Symptom Severity | **Mean (SD, range)**  28.25 (9.65, 8-54)  5.82 (2.88, 0-12)  3.37 (2.06, 0-7)  11.05 (4.85, 3-23)  8.26 (3.27, 2-15)  5.39 (2.74, 0-12) | **Mean (SD, range)**  29.93 (9.72, 8-47)  6.85 (3.30, 0-13)  3.41 (2.27, 0-8)  10.81 (5.02, 2-23)  8.85 (3.24, 4-15)  5.44 (2.24, 0-9) | **F-value**  .47  1.81  .01  .04  .52  .01 |
| ^1^Missing data on one participant in the VBFM group | | | |

| **Supplemental Table 3. Average connectivity values in regions of interest (bilateral amygdala seed) in entire sample at voxelwise p<.05 with familywise error correction** | | |
| --- | --- | --- |
|  | **Pre-intervention Mean (SD)** | **Post-intervention Mean (SD)** |
| Left hippocampus | .39 (.13) | .39 (.11) |
| Right hippocampus | .41 (.14) | .40 (.11) |
| Left insula | .06 (.13) | .04 (.11)* |

*no significant changes in connectivity observed at specified statistical threshold

| Supplemental Table 4. Scanner descriptions and acquisition parameters | | | |
| --- | --- | --- | --- |
|  | Scanner 1 | Scanner 2 | Scanner 3 |
| Scanner type | Siemens 3T Prisma_fit | Siemens 3T TrioTim | Siemens 3T Prisma_fit |
| Software | syngo MR E11 | syngo MR B17 | syngo MR E11 |
| Head Coil | HeadNeck_20 Coil | HeadMatrix Coil | HeadNeck_20 Coil |
| Number of subjects | n=5 | n=8 | n=18 |
| Modality |  |  |  |
| T1-weighted | **TR** = 2600ms, **TE** = 3.02ms **TI** = 900ms **Flip angle** = 8 deg **FOV** = 256mm **Slices** = 176 **Voxel size** = 1.0mm x 1.0mm x 1.0mm | **TR** = 2600ms, **TE** = 3.02ms **TI** = 900ms **Flip angle** = 8 deg **FOV** = 256mm **Slices** = 176 **Voxel size** = 1.0mm x 1.0mm x 1.0mm | **TR** = 2530ms, **TEs** = 1.69/3.55/5.41/7.27ms **TI** = 900ms **Flip angle** = 8 deg **FOV** = 256mm **Slices** = 176 **Voxel size** = 1.0mm x 1.0mm x 1.0mm |
| Functional MRI | **TR** = 2500ms, **TE** = 30ms, **flip angle** = 90 deg, **FOV** = 220mm, **slices** = 40, **Voxel size** = 3mm x 3mm x 3mm | **TR** = 2500ms, **TE** = 30ms, **flip angle** = 90 deg, **FOV** = 220mm, **slices** = 40, **Voxel size** = 3mm x 3mm x 3mm | **TR** = 2500ms, **TE** = 30ms, **flip angle** = 90 deg, **FOV** = 220mm, **slices** = 39, **Voxel size** = 3mm x 3mm x 3mm |

| *Supplemental Table 5* | | | | | | | | | |
| --- | --- | --- | --- | --- | --- | --- | --- | --- | --- |
|  | 1 | 2 | 3 | 4 | 5 | 6 | 7 | 8 | 9 |
| 1. MAIA Change |  |  |  |  |  |  |  |  |  |
| 2. MDI Change | -.438** |  |  |  |  |  |  |  |  |
| 3. CPT Change | .061 | -.002 |  |  |  |  |  |  |  |
| 4. LNB 0-Back Change | -.014 | .061 | .102 |  |  |  |  |  |  |
| 5. LNB 1-Back Change | -.130 | .003 | .026 | .684** |  |  |  |  |  |
| 6. LNB 2-Back Change | -.051 | -.099 | .135 | .326* | .027 |  |  |  |  |
| 7. HRV Change | .154 | -.171 | -.064 | -.011 | .126 | .328 |  |  |  |
| 8. Hippocampus-Amygdala Change to Neutral Trials | .221 | -.161 | -.066 | .180 | .163 | .019 | -.257 |  |  |
| 9. Hippocampus-Amygdala Change to Aversive Trials | .223 | -.164 | -.052 | .189 | .171 | .042 | -.240 | .998** |  |
| *p<0.05 | | | | | | | | | |
| **p<0.01 | | | | | | | | | |
| MAIA- Multidimensional Assessment of Interoceptive Awareness | | | | | | | | | |
| MDI- Multiscale Dissociation Inventory | | | | | | | | | |
| CPT- Continuous Performance Test | | | | | | | | | |
| LNB- Letter-N-Back | | | | | | | | | |
| HRV- Heart Rate Variability | | | | | | | | | |
